# Beyond the Surface: Revealing the Depths of Brain Activity by Predicting fMRI from EEG with Deep Learning

**DOI:** 10.1101/2024.11.20.624528

**Authors:** Ilia Semenkov, Pavel Rudych, Alex Ossadtchi

## Abstract

Electroencephalography (EEG) and functional magnetic resonance imaging (fMRI) are the two most commonly used non-invasive methods for studying brain function, having different but complementary strengths: high temporal resolution of the former and high spatial resolution of the latter. Crucially, fMRI is vital for studying subcortical areas, as those are practically out of reach for conventional EEG. At the same time, EEG is cost-effective and, thus, often preferable to fMRI if comparable information could be extracted which is not the case then the deep subcortical brain activity is of interest. Here we explore the possibility of recovering subcortical hemodynamics from the non-invasively recorded scalp EEG signals. To this end, we have developed a lightweight EEG-to-fMRI neural network and using an extended and publicly available dataset with concurrently recorded EEG-fMRI data show that our model allows for the prediction of the detailed Blood Oxygenation Level Dependent (BOLD) activity of 7 bilaterally symmetric subcortical structures solely from multichannel EEG data. We report the performance significantly above chance and exceeding the scores achieved for a single subcortical structure and obtained on the proprietary datasets. In contrast to the studies focusing on a single subcortical region in our approach we were able to decode multichannel EEG into 14 + 4 region-specific variations of BOLD signals measured relative to their mean hemodynamic activity. The use of relative BOLD signals allowed us to exert control over the artificial inflation of decoding accuracy scores when the decoder predicts the common mode component that is likely to have a non-neuronal origin (heartbeat, movement, etc). Finally, we interpreted our model. The electrical activity of the sensorimotor cortex appeared to contribute most to the prediction of the subcortical hemodynamics. Also, the hemodynamics of the thalamus has the smallest delay with respect to the EEG signals. Both observations are physiologically plausible and ensure the potential reliability of the decoder. Taken together, these findings pave the road towards the creation of low-cost AI-powered EEG-based fMRI digital twin technology capable of tracking subcortical activity in an ecological setting. The technology, once mature, will find numerous applications from fundamental neuroscience through diagnostics to neurorehabilitation and affective neurointerfaces.^1,2^

## 1 Introduction

Electroencephalography (EEG) and functional magnetic resonance imaging (fMRI) stand out as the top choices among non-invasive techniques for gauging brain activity, each presenting distinct advantages and drawbacks. While EEG captures the collective activity of widespread neural populations, its interpretability is somewhat constrained. Nevertheless, its unique ability to capture brain processes with millisecond precision offers valuable insights into swift cortical dynamics. Conversely, fMRI, which detects changes in Blood Oxygenation Level Dependent (BOLD) signals, boasts superior spatial resolution compared to EEG. However, its reliance on the hemodynamic responses makes this modality slow, with characteristic response time on the order of a second when scanning the entire brain. Moreover, the disparities between EEG and fMRI extend beyond the fundamental aspects. EEG devices are affordable, portable, and suitable for routine home use, while fMRI machines are cumbersome, and costly and record the data in a highly non-ecological setting.

However, despite these notable limitations, the fMRI technology remains the modality with an unparalleled capability to reliably capture the activity of deep cortical regions (such as the hippocampus) and subcortical structures (like the basal ganglia) in a non-invasive manner. The limited accessibility of the fMRI technology contributes to the historical underrepresentation of human subcortical regions in functional brain studies [1]. Yet, it is hard to overestimate the significance of these regions in cognitive processes—such as memory, attention, and reward mechanisms [2], motor functions [3], and affective cognition [4]. Moreover, subcortical structures are increasingly implicated as key factors in various neuropsychiatric disorders due to the deficiencies in both structure [5] and function [6] associated with a disease. Consequently, there is an urgent need for accessible, cost-effective, and non-invasive tools to visualize the functioning of subcortical brain regions, including the basal ganglia, cerebellum, and thalamus.

One strategy for getting access to the workings of the *subcortical brain regions* involves augmenting the accessible and reasonably ecological EEG with a deep learning model capable of predicting subcortical BOLD signals solely from the electrical signals measured with EEG. One can see the outline of this approach in Figure 1

**Figure 1:**
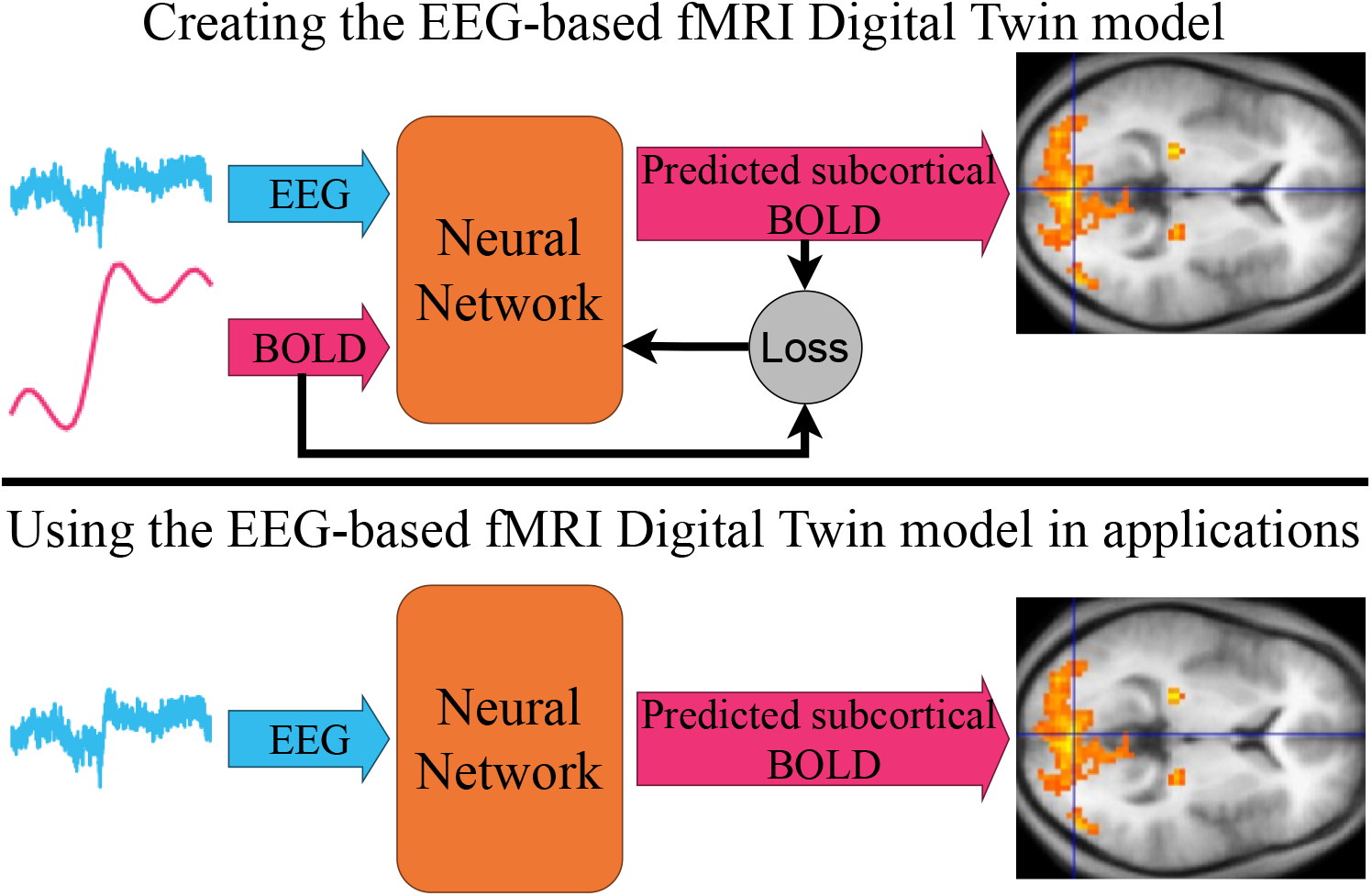
EEG-based fMRI digital twin principle. Concurrently recorded EEG-fMRI data are used to train a *subject-specific* neural network to predict regional BOLD signals from the multichannel EEG data. Once the model is built, EEG alone passes through the model which predicts the brain’s hemodynamics.

Many studies have sought to validate the congruence between EEG and fMRI findings regarding underlying brain activity by directly correlating the two modalities. To synchronize EEG and BOLD signals, a convolutional transformation is typically applied to the EEG-derived source power profiles, as demonstrated by Martinez et al. [7] and Laufs et al. [8]. An intriguing heuristic was proposed in [9], suggesting a nonlinear model that posits a relationship between the increase of BOLD signals and the broadening (expansion) of EEG signal spectra. While much of this research has concentrated on the cortical BOLD activity [10], efforts to recover subcortical BOLD (sBOLD) signals from EEG data remain a struggle.

Nonetheless, Singer et al. [11] recently presented a notable exception, demonstrating the prediction of striatum BOLD activity from EEG data in a task-based context. Notably, the authors used a simple partial least squares (PLS) model that achieved prediction accuracy based on the heuristically selected 10 EEG channels and achieved a 0.26 correlation coefficient between the ground truth and EEG-derived striatum BOLD signal. This important study, to the best of the authors’ knowledge, remains so far the only work dedicated to EEG-based subcortical BOLD activity prediction. Although significant, it focused only on the single brain region BOLD signal. This may lead to overestimated metrics due to the presence of a common mode component in the ground truth BOLD data, which can be readily predicted from the ongoing EEG and mask the individual BOLD signal variations pertinent to this specific brain structure. Moreover, of primary interest to fundamental neuroscience and clinical diagnostics is the temporal dynamics of the spatial distribution of BOLD signals across multiple brain structures [12].

To further our understanding of the EEG-BOLD relationships and aiming towards developing the EEG-based fMRI digital twin technology we are presenting our exploratory solution capable of simultaneous prediction of BOLD activity of several subcortical structures using concurrently recorded multichannel EEG. Simultaneous recovery of BOLD activity of several deep brain regions allows us **to exert control over the inflation of decoding accuracy metrics** that may potentially arise due to the common-mode component. To this end, we focus on the prediction of the relative (differential) regional BOLD activity signal obtained from the BOLD time series by subtracting from it the BOLD signal representing the average activation of all regions of interest. We selected 7 bilaterally symmetric (total 14) subcortical ROIs and also included two bilaterally symmetric hippocampi and the averaged cortical hemispheres as the additional ROIs which resulted in a total of 18 anatomically defined brain structures simultaneously predicted from the multichannel EEG time series. To the best of our knowledge, **this is the first demonstration of a successful simultaneous recovery of BOLD signals reflecting the relative hemodynamics of multiple subcortical nuclei and the hippocampus**. Also, the presented work **is the first deep learning study** using the publicly available naturalistic viewing fMRI+EEG dataset [13] and aimed at predicting **the *relative* BOLD activity of subcortical** brain regions from the concurrently recorded cleaned EEG time series. Although we could not obtain the data described in [11] we have implemented their version of the mixed norm PLS, verified our implementation on a simulated dataset, and applied it to the publicly available data EEG-fMRI data used here [13] as a baseline solution. Finally, by **providing public access to the code of our model, the baseline solution, and the entire pipeline along with the updated version of the cleaned EEG data meticulously synchronized with** the **BOLD time series**, we aim to establish a platform for streamlining both basic and applied research endeavors focused on creating tools for accessible and ecologically valid non-invasive imaging of subcortical brain activity. Once mature, the developed EEG-based fMRI digital twin technology will not only advance future studies in affective, memory, and decision-making neuroscience but also contribute to the identification of biomarkers for neurodegenerative diseases rooted in abnormal subcortical brain activity.

## 2 Dataset

The original dataset described in [13] includes simultaneously collected recordings from 22 individuals (ages: 23–51) across various visual and naturalistic stimuli. **To the best of our knowledge, this is the largest publicly available and documented dataset containing concurrently acquired raw EEG and fMRI data**. The EEG-fMRI data were collected at rest and during several visual tasks including flickering checkerboard, the visual paradigm Inscapes, and several short video movies representing naturalistic stimuli. The EEG was recorded using an MRI-compatible system with 61 cortical channels at a 5 KHz sampling rate with channel impedance values kept below 20 KOm. The functional MRI signal was registered using a 3T Siemens TrioTim equipped with a 12-channel head coil. BOLD fMRI sequences were acquired with TR = 2.1 s per volume that comprised 38 slices with planar dimensions 64 × 64 voxels of dimension [3.469 × 3.469 × 3.330] mm.

For our analysis, we have selected the EEG-fMRI recordings of the 16 subjects who appeared to have the complete set of recordings with two sessions per condition. We have excluded several subjects due to excessive movements which became apparent during the pre-processing of the raw data. Therefore our sample included 11 subjects. The total duration of data per subject was approximately equal to 7200 seconds.

## 3 Model

The majority of existing architectures for analysis of EEG data use explicit trainable, not separated by non-linearity spatial and temporal convolution layers followed by more or less standard convolutional and/or fully connected layers with non-linear activation functions in between. Such a processing is physiologically plausible as the authors typically state and accepted in the EEG community processing flow which aligns well with well-established classical approaches for the analysis of multichannel electrophysiological signals [14, 15]. The spatial filter focuses on a specific neuronal source and the temporal filter isolates the brain rhythms of interest. When this combination of spatial and temporal filters is followed by a non-linearity and subsequent smoothing one gains access to the envelope of a narrow band process reflecting the activity of a specific neuronal population generating the specified rhythm. There is currently a consensus in the EEG and MEG-based functional neuroimaging community that the electromagnetic forward modeling based on forward operator computed by solving Maxwell equations within a volume conductor (the head) accurately reconstructs non-invasively measured EEG signals produced by a cohort of neuronal sources. Due to the linearity of Maxwell equations, it is conceivable that the unmixing process would also rely on a linear operation such as spatial filtering that prescribes computing a linear combination of sensor signals with the spatial filter weights.

At the same time, things are significantly less straightforward with EEG’s temporal structure whose analysis may require significantly non-linear processing even at the early feature extraction step. While indeed rhythmic activity plays a pivotal role in communication between neuronal assemblies [16] and hallmarks resting state of specific neuronal populations, recent studies also demonstrate a growing interest in the aperiodic component of the electrophysiological signals [17, 18, 19].

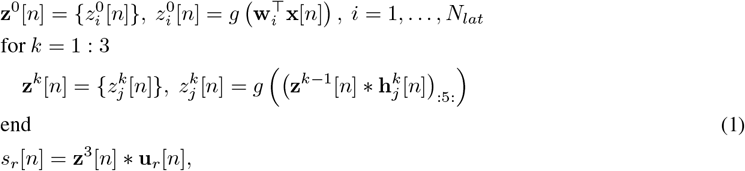

where *j* = 1, .., *N*_*lat*_, *k* = 1, .., 3, *r* = 1, .., *N*_*lat*_, **w**_*i*_ are [*N*_*EEG*_ *×* 1] spatial filter vectors or 1D convolution kernels, 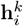 are [*N*_*lat*_ *×* 5] 1D strided convolution kernels with stride 5 as denoted by (.) _:5:,_, **u**_*i*_ are the [*N*_*lat*_ *×* 21] 1D convolution kernels, *g*(.) is the GELU activation function, and **a**[*n*] * **b**[*n*] denote the 1D convolution operation where for each position of the filter the overlapping temporal data samples are scaled with the filter weights for all channels and then lumped together. Note that for clarity and compactness we excluded the bias terms in these expressions.

In our architecture, see Figure 2 and equations (1), we use only linear spatial filtering and convolve over the temporal dimension the already non-linearly processed data. The spatial filtering, performed by a convolutional layer with a kernel size of 1, maps *N*_*EEG*_ = 61 EEG channels into *N*_*lat*_ = 23 latent channels. Afterward, we apply the GELU activation function. Then we pass the resultant time series to the sequence of 3 Pyramidal subsampling modules. Each of these modules comprises a convolutional layer with both stride and kernel size equal to 5 samples followed by the GELU activation and dropout with 0.25 probability to avoid overfitting. This sequence of layers not only downsamples the 250 Hz EEG signal to the desired 2 Hz of the BOLD but also acts as a series of temporal filters with increasing kernels. The name Pyramidal subsampling is inspired by [20] because each module deals with a smaller sampling rate signal while maintaining the same kernel size in samples. Therefore, a fixed kernel of 5 samples gradually expands if measured in seconds. Finally, we apply the main temporal filter of size 10 seconds + 1 sample (to keep the kernel size odd) which gives the desired BOLD signal approximations. The amount of channels is maintained at *N*_*lat*_ = 23 throughout the whole network after the initial spatial filter. It is important to note that given a nearly linear segment of the GELU function corresponding to the positive argument values and the limited magnitude of the input signals the network could still implement linear temporal filtering by appropriately adjusting bias weights present in every layer.

**Figure 2:**
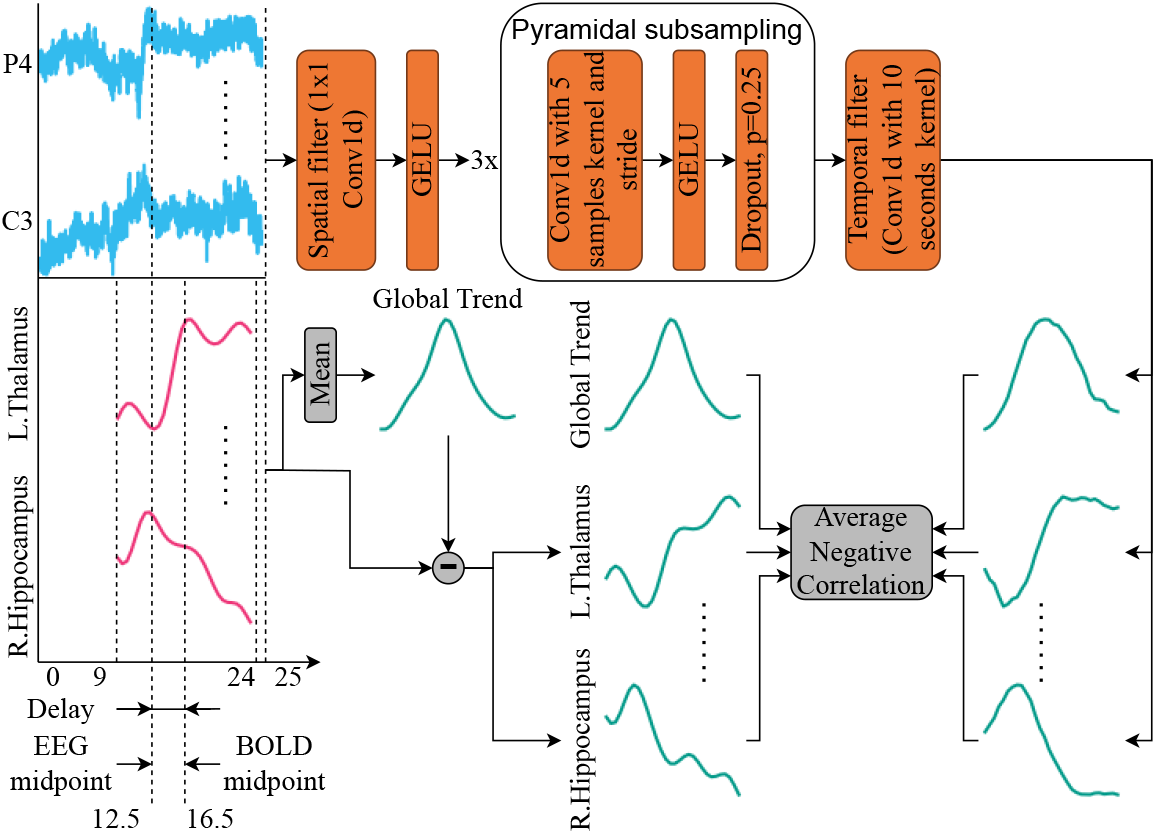
Our neural network architecture and the processing pipeline. We take an EEG sequence to be 10 seconds longer to avoid edge effects from padding in the temporal filter layer. One can see that BOLD sequence is delayed by 4 seconds by looking at the difference between the midpoints.

It is commonly accepted in the neurophysiological community that brain’s hemodynamic response is delayed with respect to the stimulus by 3-7 seconds for the majority of the brain structures depending on the properties of the stimulus [21]. This is reflected in the shape of the hemodynamic response function (HRF) that is frequently used to align the brain’s electrical activity and its hemodynamics by computing the convolution of non-linearly transformed electrical activity with the HRF kernel. In our model we focused its receptive field within the window, delayed with respect to the segment where the BOLD signal is to be predicted. For the majority of our experiments, we used the delay *d* = 4 seconds. The alignment between the input EEG time series is illustrated in Figure 2. We also take into consideration the transients associated with the convolution operation. To avoid edge effects during temporal filtering in the final layer of the network we use no padding and take an input segment to be 10 seconds longer. This results into having 25-seconds long input EEG segments and 15-seconds long corresponding BOLD ground truth segments, see the bottom left part of Figure 2.

Once the model was trained, we explored the spatial filter weights. It has been demonstrated earlier [15, 22] that the spatial filter weights alone contain not only information about the target neuronal sources but are also influenced by the spatial distribution of the interfering activity. This means that the spatial filter weights **w**_*i*_ need to be converted to the spatial patterns before the judgments can be made about the underlying brain sources. Therefore we have transformed the spatial filter weights **w**_*i*_ into the spatial patterns as **p**_*i*_ = **Rw**_*i*_, where **R** is the spatial covariance matrix of the EEG data estimated from the training segment of our data.

## 4 Performance metrics

As a quality metric, we used the Pearson correlation coefficient reflecting the similarity between the original and the EEG-derived BOLD activity profiles. This measure is appropriate since our main interest is to recover the fluctuations of the BOLD signal around its global mean that happen over time and are specific to each of the chosen ROIs. Also, this metric was used in the study [11], the only paper to date demonstrating the prediction of subcortical hemodynamic activity from EEG data and where the prediction performance was carefully quantified.

The EEG-fMRI data [13] were collected during both: the resting state and during the periods when a volunteer watched checkerboard and 7 videos, each corresponding to the experimental condition. When calculating the performance metrics we treated the data as the ongoing BOLD signal and did not use condition labels to aggregate the obtained brain responses within each of the conditions. This resulted in generally low scores simply because the BOLD data are known to be quite noisy. We did so because we wanted to avoid the data leak. A network could simply learn to decode from EEG the condition label and assign the output signal depending on this label. This would not correspond to our goal of learning a model capable of translating scalp EEG to the subcortical hemodynamic activity.

## 5 Training and testing strategy

The dataset contains EEG-fMRI data recorded during different tasks and the resting state [13]. For some of the conditions, the dataset contained recordings made during the two separate sessions. The total duration of the dataset per subject was over 7300 seconds or over 2 hours. Each run of duration between 200 and 600 seconds depending on the stimuli (see [13] for more details) was split into three **contiguous train-validation-test** segments in proportion 55 % - 22.5% - 22.5% and all training segments from all the runs were then lumped together and used for training. This resulted in a total of around 4000 seconds of training data per subject. This strategy as **opposed to the random shuffling** allowed us to avoid the artificial inflation of the decoding scores due to the natural temporal smoothness pertinent to the EEG and fMRI data. Using this data we trained individual models, see Section 3 for each of the subjects separately and performed testing using a total of 1650 seconds of data per subject split between the experimental conditions. These numbers may vary insignificantly as we tailored the splits aligned with fMRI volume acquisition timing with TR = 2.1 sec to avoid data loss.

## 6 Results

As described in Section 5 we have trained individual models for each of our 11 subjects to predict region-specific hemodynamics from the scalp EEG data. The averaged over all subjects decoding accuracy measured as the correlation coefficient between the original and the EEG-predicted BOLD signals, see Section 4, is shown in Figure 3a for 18 brain structures and the common mode regional global trend BOLD signal. As expected the global trend BOLD signal appears to be predicted most accurately with a significantly non-zero mean correlation coefficient of 0.44. This is exactly the reason why in this work we decided to focus on the **relative** activation of brain structures with respect to their global mean instead of trying to predict the absolute signals containing the significant common mode contribution. Not doing so would drastically boost the accuracy scores without bringing us any closer to our final goal.

**Figure 3:**
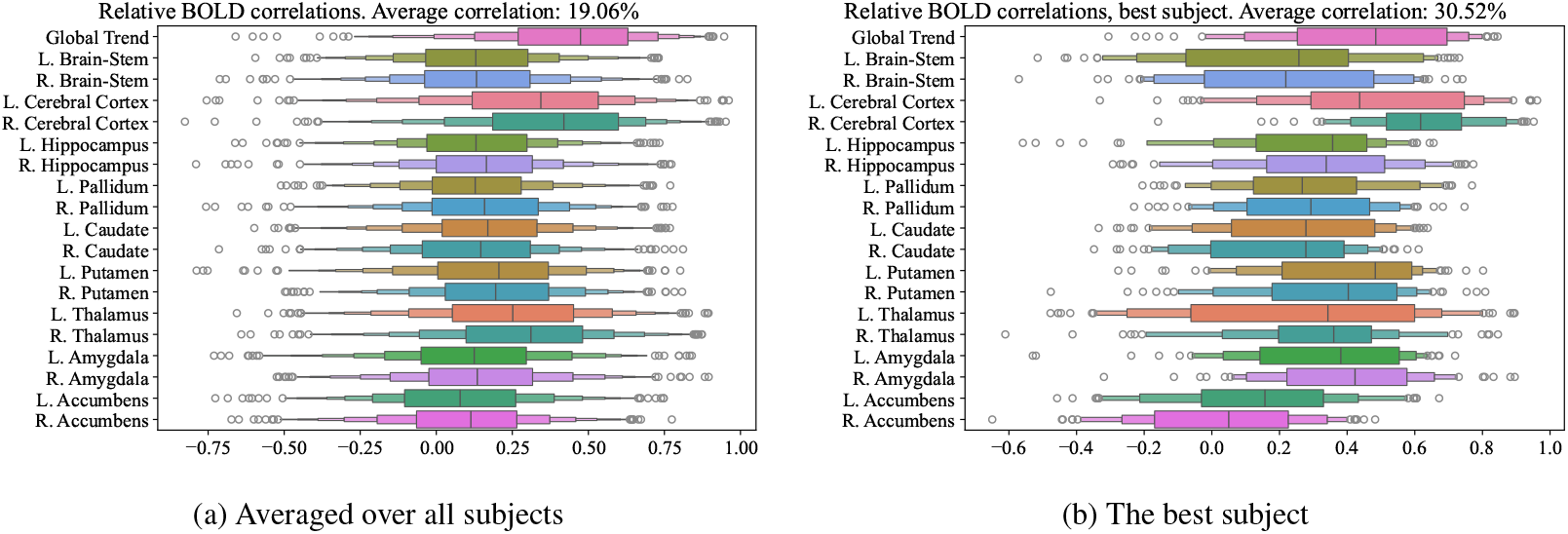
Decoding correlations of the **relative** BOLD obtained on Natural viewing publicly available dataset averaged over all tasks and runs.

Indeed, it appears much harder to predict the relative BOLD signal of the subcortical structures which is expressed by the average over all regions, runs, and subjects correlation coefficient of 0.19. At the same time, we observed large deviations in the regional correlation coefficient across subjects and runs. When interpreting these results one has to keep in mind that not all the brain structures exhibit pronounced activity during the experimental conditions represented in the dataset. For example, the low prediction performance score for nucleus accumbens hemodynamics may be explained by the fact that the resting state and the repertoire of the tasks in the Natural viewing dataset failed to activate this structure that is known to be the key node of our brain’s reward processing system and theoretically should not be activated during the passive watching of a video stream. At the same time, we are observing relatively higher scores for both the left and right thalamus, the subcortical brain structures known to modulate the default mode network activity [23, 24]. The contribution from the latter was shown to dominate during the movie watching [25]. Relatively high scores for bilateral putamen and caudate are in line with the observation that these two are activated during video perception and are involved in the visual attention mechanism maintenance [26].

The corresponding plot showing regional prediction accuracy distributions for the best-performing subject is shown in Figure 3b. The model appears to predict the relative BOLD signal with an average correlation of 30%. The global trend signal is recovered with a correlation of 50% and the nucleus accumbens activity prediction still demonstrates the worst performance.

The current State-of-the-Art study [11] utilizes the following pipeline. Firstly, the authors transform original EEG signals into band power time series in a selected sequence of frequency bands. This procedure also downsamples the obtained band-power profiles to 2 Hz (the same sampling rate as fMRI). Then, they use an adaptation of the group LASSO approach to find a subset of up to 10 channels to be used in the regression. Finally, they apply the Partial Least Squares regression to the resultant band-power profiles in a subset of chosen channels. In the time domain, they utilize 60 lagged values with a step of 0.5 seconds to predict the BOLD signal. For more details see the original paper [11]. Because the main trainable model is the Partial Least Squares we refer to this whole pipeline as PLS.

A detailed performance comparison of our approach to the PLS can be seen in Table 1. Our model noticeably outperforms the PLS approach over almost every single ROI. The only ROIs that PLS predicts slightly better are the Right Putamen, Right Brain-Stem, and Right Accumbens. On average our model achieves a correlation of 19.06% and statistically significantly outperforms the PLS with *p* ≪ 0.01 as indicated by the one-sided Wilcoxon signed-rank test.

**Table 1:**
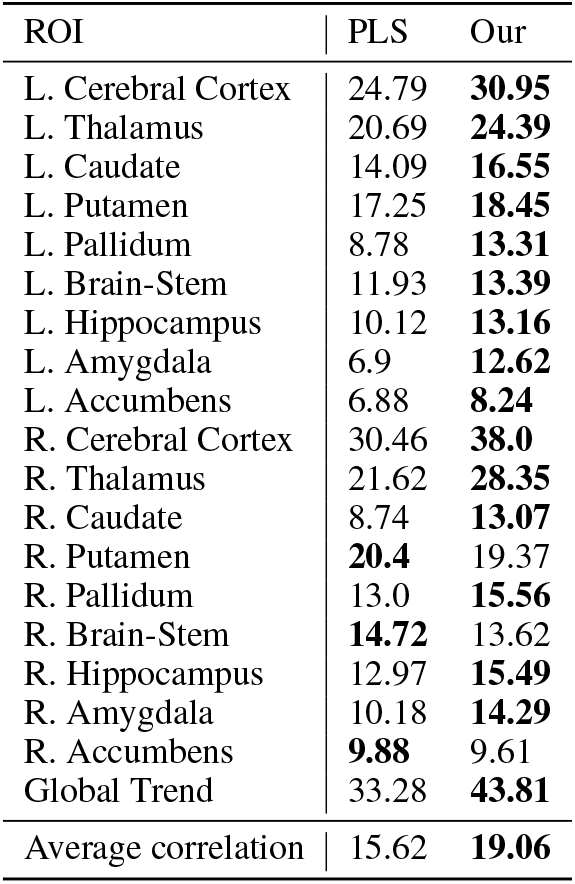
Test correlations per ROIs for Partial Least Squares pipeline (PLS) and our neural network. Correlation values are averaged across all tasks and subjects.

## 7 Interpretation

To check for the significance of the reported correlation scores we have systematically shifted the EEG data with respect to the corresponding synchronized fMRI time series which resulted in a relative shift value from -5 seconds to +10 seconds. Then, we treated these pairs of mutually shifted EEG and BOLD time series data as the new dataset, trained our model anew, and measured the average performance over five training runs for each ROI and each tested relative shift value. The results of this experiment are shown in Figure 4.

**Figure 4:**
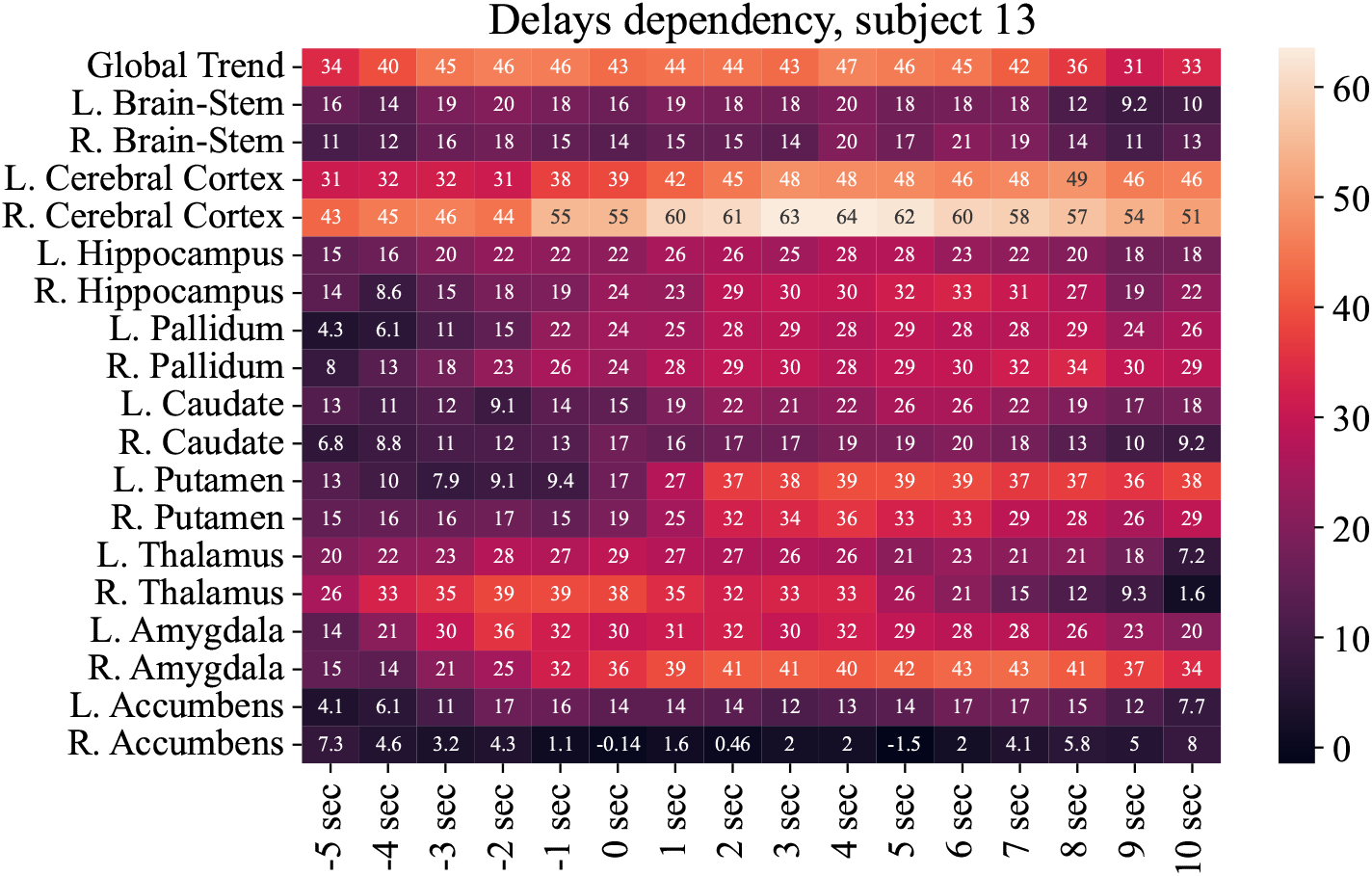
Decoding accuracy as a function of EEG vs. fMRI delay for individual brain regions. See also Figure 2 and Section 3 for the description of the EEG vs. fMRI alignment strategy.

We can observe that on average the delay of 5-6 sec appears to correspond to the best prediction scores which is consistent with the known location of the HRF’s peak. At the same time, individual ROIs exhibit certain variability in the optimal shift value. Interestingly, BOLD activity of the thalamus appears to have the smallest lag with respect to the scalp EEG which is consistent with the observation reported in [27] and may reflect the reported significantly faster hemodynamic timing in the lateral geniculate nucleus of the thalamus.

Next, in Figure 5 we present the topographic plots of the six most powerful patterns (patterns with the largest L2 norm) corresponding to the spatial filters [22] in our model, see Figure 2. The first pattern emphasizes a source in the sensory-motor region which is consistent with the functional role of basal ganglia and the presence of the corresponding anatomical links between the sub-cortex and sensory-motor cortical areas [28]. The neuronal sources corresponding to the second and third patterns are located in the parietal region and may reflect the increased attention of our model to the generators of the resting state alpha-band activity. The last three patterns correspond to either deep brain sources or may potentially reflect the tuning of our model to the cardiovascular activity which may help the model to compensate for the influence of the cardiac activity that is supposed to be minimally present in the target **relative** BOLD signals.

**Figure 5:**
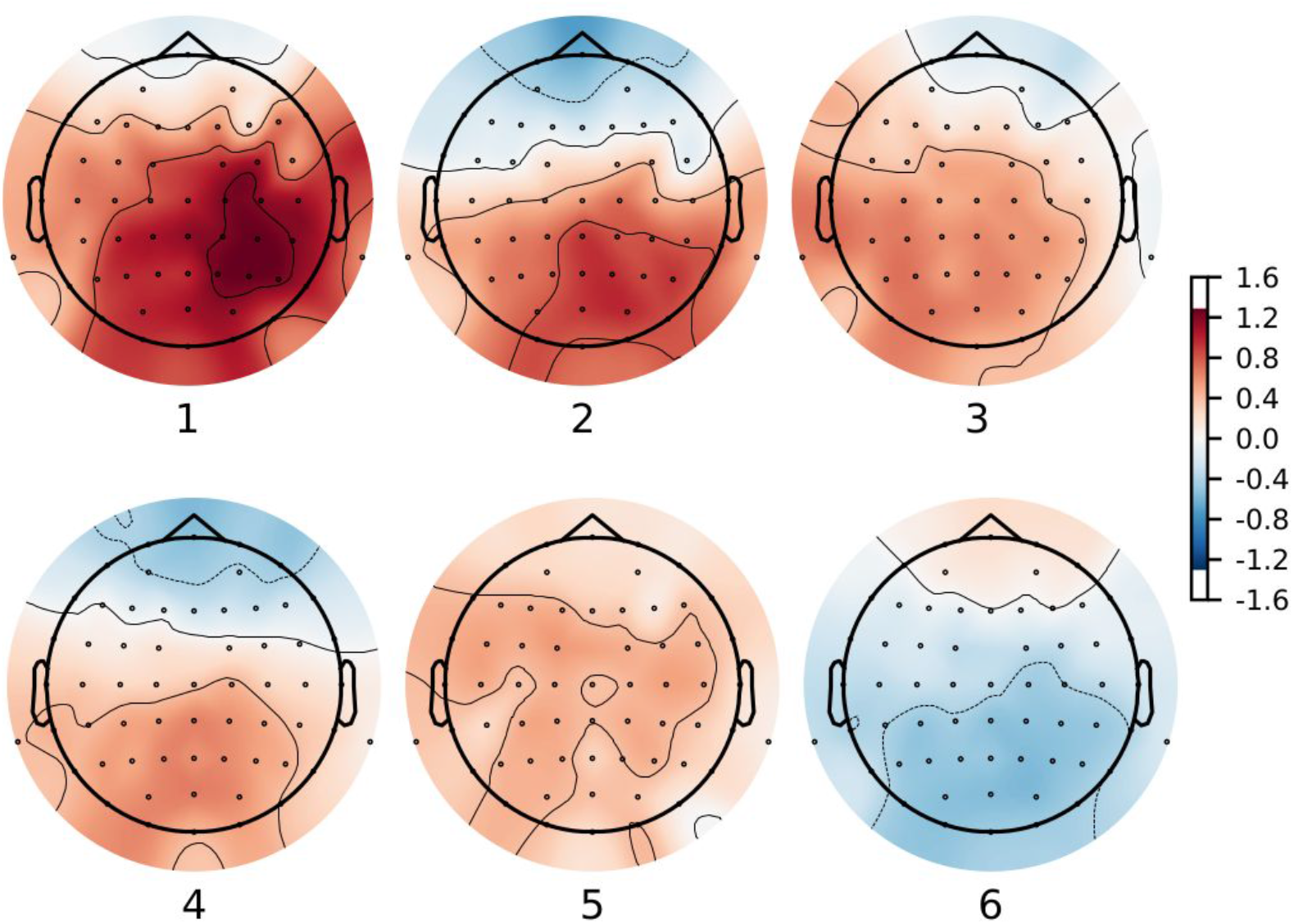
Topographic plots of the 6 most powerful patterns [22] corresponding to the spatial filters of our model.

## 8 Exploring time-series during specific conditions

In Figure 6, we demonstrate the original (derived from fMRI) and the predicted from EEG BOLD time series of relative activation for 9 bilaterally symmetric brain structures (7 pairs of subcortical regions, the pair of integrated cerebral cortices and hippocampi) in the resting state (Figure 6a) and the “Inscapes” condition (Figure 6b). During this condition, the subjects were viewing a 3D computer-generated animation featuring abstract 3D shapes and moves in slow continuous transitions. We can observe reasonable agreement between the original ground truth (black) and predicted (yellow) traces. Interestingly, while in the resting state condition brainstem’s BOLD gets predicted well, the model fails to track the brainstem’s hemodynamics during the visual task. At the same time the activity of putamen and caudate, the two structures comprising the dorsal striatum, appear to be decoded well from the scalp EEG data in both the resting state and the visual task conditions with almost two-fold increased correlation as compared to the performance reported in the current state of the art study [11]. We also note that in contrast to the existing studies we are reporting the performance for relative BOLD signals reflecting region-specific activity with global trend subtracted which exerts a reasonable control over potential inflation of decoding quality metrics due to taking into account the confounding factors of non-neuronal origin.

**Figure 6:**
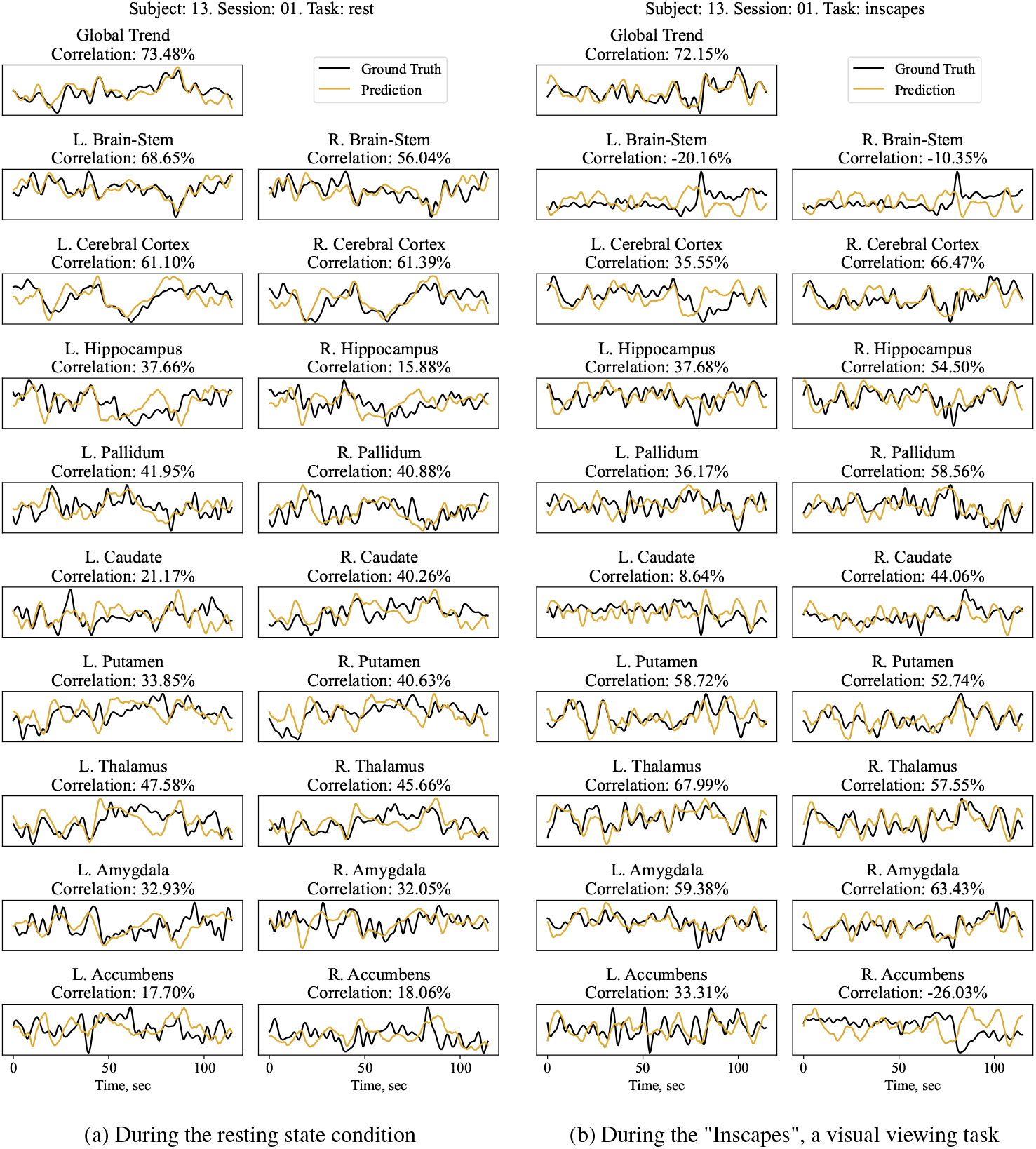
Original (black) and the EEG-derived BOLD time series for 9 bilaterally symmetric brain structures and the common mode Global trend component.

## 9 Discussion

Technological advances now allow for recording the brain’s electrical activity concurrently with BOLD signals within the fMRI scanner [29], [30]. This possibility triggered a range of studies aimed at relating fMRI and EEG signals in various experimental paradigms. The majority of attempts establish the similarity of EEG and fMRI-based findings about the underlying brain activity by directly correlating the two measures with the prior transformation of the EEG data to the fMRI scale by a simple convolution of the pre-processed EEG with Hemodynamic Response Function, e.g. [31], [32]. Intriguingly, it has been recently shown that cortical BOLD signal can be reconstructed from multichannel EEG data utilizing machine learning [33]. Leite in [34] attempted to build a transfer function between EEG and BOLD signals in the context of analysis of epileptic activity. Dominantly, the existing studies exploring the EEG-BOLD relationship, see [35], focus on the analysis of cortical activity and not the subcortical structures such as basal ganglia and other deep brain regions that appear responsible for a broad range of behaviors, reflecting emotional state, underlying complex decision making and risk-reward balancing processes, voluntary movements control, memory and various kinds of learning.

Last year’s work [11] described an interesting attempt to build a simple partial least squares-based solution to recover the BOLD signal of the ventral striatum, the key subcortical region responsible for reward processing and anticipation. The authors achieved *r* = 0.21 correlation coefficient between the ground truth and the EEG-derived BOLD and replicated the result with *r* = 0.26 on the out-of-sample dataset. Although this impressive result has been state-of-the-art, the observed performance metrics may be potentially misleading since as the authors discuss, the reported scores may result from the presence of the common mode component in the regional striatum signal which may potentially inflate the obtained performance characteristic. To illustrate this claim in Figure 7 we show the average prediction accuracy of the absolute (not relative) BOLD achieved by our model on the naturalistic viewing dataset at hand. Comparing this diagram with the one presented in Figure 3a we can notice more than 1.5 fold inflation (33.6% vs. 19.06%) of the average over the tasks, subjects and runs decoding accuracy explained purely by the lack of removing the common mode component from the ground truth regional ROI time series. Unfortunately, we were not able to obtain the dataset used in that study but we did our best to implement the described approach and used it in our comparative performance analysis, see Table 1.

**Figure 7:**
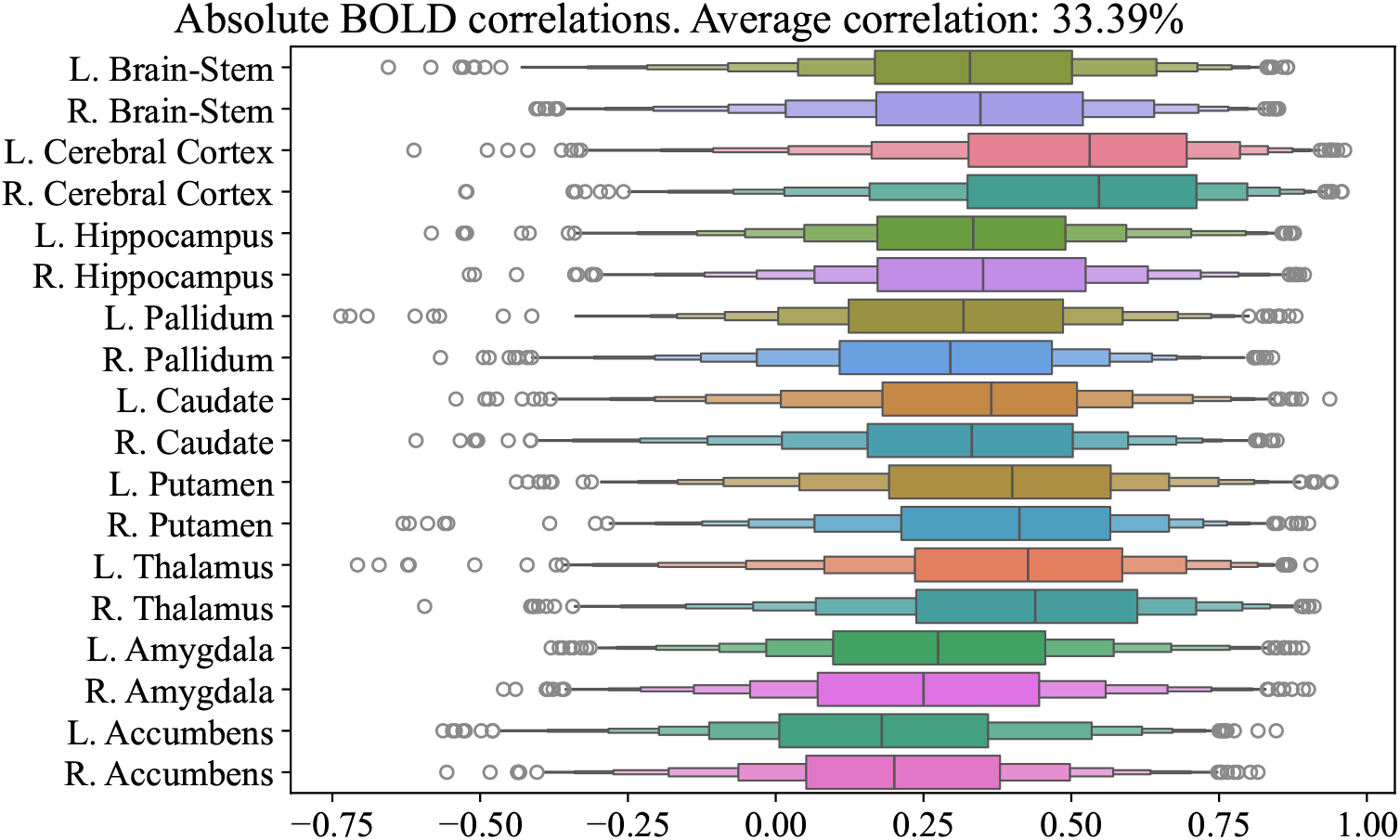
Decoding accuracy of **absolute** BOLD averaged over all tasks and subjects.

Also, note that this work did not aim to create a general multi-subject model for two reasons. First, the amount of available data is not enough to learn the needed generalization even with the use of subject-specific layers accounting for subject-specific variations in the brain’s anatomy and the EEG electrodes placement.

Second, we assume that in the early stages, the EEG-based fMRI digital twin technology will not only require individual models, but will use a subject-specific model built immediately before the EEG-only experiment. This will nevertheless represent a big leap forward furnishing the opportunity for ecological exploration of subcortical activity in the cognitive, decision making and motor experimental paradigms.

## 10 Limitations

Although we have used the most comprehensive and extended publicly available EEG-fMRI dataset, one of the key limitations of this work is the relatively small amount of data which precluded us from building and training more sophisticated models capable of generalizing over all subjects and conditions. We, therefore, resorted to building subject-specific models that nevertheless appear relevant within the expected use of the developed EEG-based fMRI digital twin technology where each subject has one’s own model to convert scalp EEG to the subcortical hemodynamics. Limited amount of data can be ameliorated in the future by using encoder-decoder architectures pre-trained on a large amount of openly available unimodal EEG and fMRI datasets.

The decoding accuracy can be improved by using cleaner EEG signals. The absolute majority of the existing concurrently collected EEG+fMRI data such as the dataset used here [13]are recorded in the paradigm where EEG and BOLD acquisition is performed continuously. This leads to contamination of the EEG signal with very strong high-frequency gradient artifacts. However, it is known that the peak of the hemodynamic response function [36] is centered at around 5-6 seconds, our observations, see Figure 4 also support this claim but show interesting variations in the HRF latency depending on the brain regions of interest. These theoretical and empirically observed lags suggest the following intermittent EEG+fMRI acquisition paradigm. In this paradigm, when recording EEG to be used for prediction of the forthcoming BOLD signal we will switch the gradient coils off for 4-6 seconds to yield broadband artifact-free EEG. Then, by turning the gradient coils on we will collect several volumes of the BOLD signal to be predicted from the preceding EEG data. Then the coils are switched off again and the cycle repeats. This paradigm will produce cleaner and more informative EEG data which is likely to improve the BOLD reconstruction accuracy. An additional benefit here is that the EEG data collected in such an intermittent paradigm will more resemble the signals observed outside the scanner and used in the inference mode of the EEG-based fMRI digital twin technology once it matures, see Figure 1. An additional strategy for obtaining cleaner EEG data recorded within an fMRI scanner lies in using carbon wire loop technology [37].

## A Appendix

### A.1 fMRI pre-processing

fMRI data in NIFTI format was aggregated in the BIDS folder and pre-processed with fMRIPrep [38] pipeline with automatic removal of ICA Components classified by AROMA and smoothing. Confound regression with respect to the motion and global signals including those from the white matter as regressors was used to remove from BOLD activity the signals of non-neuronal origin. During the pre-processing the fMRI data was normalized to the MNI space, slice-timing corrected, and smoothed with a spatial Gaussian filter (5-mm kernel).

Subsequently, we extracted the BOLD activity of bilaterally symmetric regions of interest (ROI) using the Harvard-Oxford structural atlas [39]. We focused on the list of 14 bilaterally symmetric subcortical ROIs comprising nucleus accumbens, amygdala, caudate, pallidum, putamen, thalamus, and two lateralized brainstem ROIs. We have also included two bilaterally symmetric hippocampi and the averaged cortical hemispheres as additional ROIs.

Importantly, to avoid the influence of the common mode signal on the obtained performance we computed the averaged over 18 regions of interest (ROI) BOLD signal time series and subtracted it from the individual ROI BOLD time series. **This allowed us to focus on decoding the ROI-specific fluctuations of the functional hemodynamics (relative regional BOLD signal) and to avoid the artificial inflation of the obtained performance scores by the factors of non-neuronal origin**. The averaged over ROIs BOLD signal was then treated as a separate ROI dubbed ‘Global trend’.

### A.2 EEG data pre-processing

We used raw EEG data provided in the dataset as they represented the most straightforward alignment with the BOLD time series. The raw EEG was cleaned from the broadband gradient [40] and cardioballistic artifacts [41] using BrainProducts Analyzer software. The rest of the fMRI influence was removed using ICA followed by a manual component classification based on the visual analysis of their spatial and spectral properties. We also monitored the power spectral density of the resultant time series to guide this iterative process.

### A.3 Training details

The model was trained using the average over batch and ROIs negative correlation as the loss function. We also used the Adam optimizer [42] with the ReduceLROnPlateau scheduler which divided the learning rate by 10 if validation loss did not improve over 5 consecutive epochs. We also utilized early stopping which would terminate training after 20 epochs without validation loss improvements to combat overfitting.

Each training run for a single subject took about 20 minutes (depending on early stopping) using a single NVIDIA V100 GPU with 32 GB VRAM. Used libraries and frameworks could be found in the requirements file in the provided repository.

Because the BOLD signal is known to be out of sync with the EEG signal, we were trying different delay values as a hyperparameter. The range which we tried is from -5 seconds (BOLD before EEG) and up to 10 seconds (BOLD after EEG). We ultimately chosen delay equal to 4 seconds which aligns with the theoretical results on the topic. One can see the tested range at Figure 4. Additionally, we tested ReLU as an alternative activation function, but ultimately went with GELU. Also, we varied the dropout probability in every Pyramidal subsampling layer from 0 (no dropout) to 0.4. The final choice landed on 0.25. All of these choices were made based on the validation metrics. The model used for the experiments is the one with the best validation correlation. All the details and configuration files of the final model are available in our repository.

For the PLS pipeline we utilized the recommended grid search ranges from the original paper [11]. We scanned through number of components for the PLS ranging from 1 to 10. And the number of the most important EEG channels also went from 1 to 10. The best combination of parameters was chosen using the grid search correlation values.

## Footnotes

1 Our dataset is available here: https://zenodo.org/records/11246524

2 Our code is available here: https://github.com/AIRI-Institute/EEG-BOLD-Decoding

